# EGFR-Mediated Mechanotransduction in Aortic Valve Cells: A Key Pathway in Response to Wall Shear Stress

**DOI:** 10.1101/2025.03.10.642402

**Authors:** Amélie Gasté, Damien Marchese, Adèle Faucherre, Louna Lopez, Eric Bertrand, Alexis Théron, Marine Herbane, Amel Seddik, Chris Jopling, Jean-François Avierinos, Hafid Ait-Oufella, Valérie Deplano, Stéphane Zaffran

**Author notes:** These authors contributed equally to this work (Amélie Gasté, Damien Marchese).

## Abstract

**Aim:** Blood flow-induced mechanical forces, particularly wall shear stress (WSS), play a fundamental role in aortic valve remodeling and maturation. Dysregulation of these processes contributes to age-related valve diseases, such as aortic stenosis and regurgitation. While epidermal growth factor receptor (EGFR) signaling has been implicated in valve development, its role in mechanotransduction remains unclear. This study aims to investigate how EGFR regulates WSS-induced signaling in valvular cells and explore its interaction with the mechanosensitive ion channel PIEZO1.

**Methods and Results:** To investigate the role of EGFR in valvular cell mechanotransduction, we used conditional *Egfr^flox^* allele to selectively delete *Egfr* in valvular cells. Histological analysis revealed increased valve leaflet thickness and hyperproliferation of mesenchymal cells when Egfr was deleted both endothelial (Tie2-Cre lineage) and mesenchymal (Sm22α-Cre lineage) cells. This was accompanied by a reduction in maturation-related genes (*Egr1, Nos3, Tgf-β*) and extracellular matrix (ECM) components. We previously demonstrated that *Egr1* expression is regulated by WSS in valvular endothelial cells, prompting further exploration of Egr1’s role in valvular cells. In vitro, Egr1 overexpression and shRNA-mediated knockdown confirmed its role in regulating *Nos3, Col1a1, and Tgf-β*, key mediators of valve remodeling. Using a pulsatile WSS-mimicking device, we found that WSS induces Erk1/2 phosphorylation and Egr1 expression in valvular cells, both of which were abolished by EGFR inhibition. However, direct EGFR activation via EGF failed to replicate WSS-induced *Egr1* expression, suggesting the involvement of additional mechanosensitive pathways. Pharmacological studies further revealed that PIEZO1 inhibition impaired WSS-induced *Egr1* expression, while PIEZO1 activation (via YODA) mimicked WSS effects on Erk1/2 phosphorylation and *Egr1* expression. These findings suggest a functional interaction between EGFR and PIEZO1 in mechanotransduction, linking mechanical forces to key molecular pathways in valve remodeling.

**Conclusion:** Our findings establish EGFR as a critical mediator of WSS-induced mechanotransduction in valve remodeling, working in synergy with PIEZO1 to regulate flow-sensitive transcription factors such as Egr1. This study provides new insights into the molecular mechanisms governing valve maturation and highlights potential therapeutic targets for age-related valve pathologies linked to abnormal WSS responses.

**Translational Perspective:** Our study highlights the pivotal role of EGFR and the mechanosensitive ion channel PIEZO1 in aortic valve cell responses to wall shear stress (WSS), offering new insights into valve remodeling. The findings suggest that dysregulated EGFR signaling contributes to valve thickening and stenosis, key factors in age-related valvular heart disease. By identifying EGFR-PIEZO1 as a critical mechanotransduction pathway, our work provides a potential therapeutic target for early intervention in aortic valve disease. Modulating EGFR or PIEZO1 activity could help mitigate pathological valve remodeling, presenting novel strategies for treating or preventing degenerative valve disorders.

## Introduction

Tissue morphogenesis can be influenced by mechanical forces, which play a pivotal role in orchestrating the cellular and molecular processes that determine cell fate ^1^. Aortic valve remodeling exemplifies this relationship by adapting its structure and function in response to the mechanical properties of the blood flow it encounters. Over time, due to mechanical stress like wall shear stress (WSS), fibrosis and calcification can appear on the valve, change its biomechanical properties and can lead to valve stenosis ^2,3^. During heart development, the aortic valve forms at the endocardial cushions within the outflow tract (OFT), where signals from the endocardium and myocardium guide endothelial cells to invade the surrounding extracellular matrix (ECM) ^4^. The WSS generated by the blood flow contributes to the proper establishment of these structures ^5,6^. As the endocardial cushions develop and enlarge, they fuse to separate the OFT into two distinct channels. The unfused margins of the septal cushions form the coronary leaflets, while the posterior intercalated cushion gives rise to the non-coronary leaflet of the aortic valve ^7^. This is followed by the valve maturation stage, during which the number of cells within the leaflets decreases, and a specialized ECM is produced with three distinct layers: the fibrosa, spongiosa, and ventricularis. Each layer has a unique protein composition specifically adapted to withstand the biomechanical stresses imposed on the valve ^3^.

The epidermal growth factor receptor (EGFR) family is essential for cardiogenesis, as mutations in the *Egfr* gene are linked to aortic valve abnormalities ^8,9^. EGFR, also known as ErbB1, is a tyrosine kinase transmembrane receptor activated by multiple ligands, including EGF, HB-EGF, TGF-α, amphiregulin, epiregulin, and beta-regulin. Ligand stimulation of the EGFR induces receptor dimerization and autophosphorylation, leading to the activation of multiple intracellular pathways including STAT, PCK, PI3K/Akt, or the MAP kinase pathway. Mutations in the *Egfr* gene have been linked to aortic valve hypertrophy ^10–13^, aortic valve disorganization, unicuspid and bicuspid aortic valves ^14,15^. During valve maturation, the activation of EGFR triggers the MAP kinase pathway and inhibits cell proliferation ^16,17^. This model has been further explored, revealing extracellular perturbations, inflammation, and calcification, highlighting its proximity to human valve stenosis ^18^.

Experimental measurements of the dynamic fluid shear stress on aortic valve leaflets reveal that the values of WSS is comprised between 46 and 70 dyne/cm² near the base and the belly of the valve leaflet to reach a maximum of 95 dyne/cm^2^ at the tip of the leaflet ^19–21^. Differences in WSS can trigger local adaptations, leading to remodeling of the ECM within the valve leaflets ^22,23^. To generate these adaptations, the valvular endothelial cells, forming the monolayer of cells around the valve leaflet, express mechanosensors such as the Piezo ion channel, VEGFR2, VE-cadherin, and PECAM ^24,25^. Their importance during valve development has been shown in zebrafish, where *Piezo1* loss of function results in defective OFT and aortic valve development ^25,26^. In response to mechanical stimuli, mechanosensors such as Piezo1, initiate a cellular response. For example, the expression of certain transcription factors, such as *Egr1* (Early Growth Response 1) and *Klf2* (Krüppel-like Factor 2), increases in response to WSS ^26–29^, and can subsequently play a role in regulating ECM composition ^30^. Several studies have shown that *Egr1* is involved in various cardiac pathologies, such as myocardial fibrosis, hypertension, pathological angiogenesis ^31^, cardiac hypertrophy, and insufficiency (related to TGF-β) ^32^.

Here, using mouse models in conjunction with the Cre-lox system, we demonstrate that EGFR plays a critical role in endothelial (Tie2-Cre lineage) and mesenchymal (Sm22α-Cre lineage) cells. Echocardiographic analysis revealed aortic valve stenosis and aortic arch dilation when *Egfr* was deleted in either lineage. Examination of the developing aortic valve showed hypertrophy driven by increased cell proliferation and decreased expression of *Col1a1*, *Tgf-β*, *Nos3*, and *Egr1*, all key factors in valve maturation. Using an *in vitro* system to apply flow activation to valvular cells, we show that EGFR signaling is crucial for the WSS response. Pharmacological manipulation revealed that EGFR signaling interacts with the Piezo1 ion channel in response to WSS, subsequently activating the MAP kinase pathway to regulate the transcription of *Egr1*, an activator of *Col1a1* and *Nos3* expression. These findings provide novel insights into the mechanistic role of EGFR signaling in WSS response, highlighting *Egr1* as a central regulator of ECM alterations in valve disease. Our study contributes to understanding valve pathology and offers potential therapeutic avenues for intervention.

## Materials and Methods

### Mouse strains

All mice were maintained according to standard laboratory conditions and all procedures were carried out under protocols approved by i) the local Animal Welfare and Ethical Committee (Comité d’éthique CE14) and ii) the national appointed ethical committee for animal experimentation (Ministere de l’Education Nationale, de l’Enseignement Supérieur et de la Recherche; Authorization #47027-2024012411177857 v3) in line with Directive 2010/63/EU of the European Parliament. The mouse strains used in our study have been described in previous lab *Egfr^f/f^*^33^, *Tagln-Cre* (*Sm22α-Cre*) or *Tie2-Cre* ^34,35^. Timed litters were collected at embryonic day (E) 10.5 to E18.5, counting evidence of a vaginal plug as E0.5. At the end of the study, all mice were euthanized by gradually increasing concentrations of CO_2_ inhalation in an induction cage. The process including filling, maintenance and emptying phases over 10 min, was conducted in accordance to the Directive 2010/63/EU of the European Parliament.

### Echocardiography

*In vivo* valve structure and function of *Tie2-Cre;Egfr^f/f^* and *Sm22α-Cre;Egfr^f/f^* mice were evaluated using a high-frequency scanner (VevoF2 from FUJIFILM ViualSonics) as previously described ^36^. Briefly, 1 month old and 3-month-old mice were anesthetized with 1-2% isofluorane inhalation and placed on a heated platform to maintain temperature during the analysis. Two-dimensional imaging was recorded with a UHF57x transducer (53297) to capture long- and short-axis projections with guided M-Mode, B-Mode and color and pulsed-wave Doppler. Doppler interrogation was performed on the arterial valve outflow in the parasternal long-axis view to assess aortic flow.

### Histological and immunofluorescence analysis

Standard histological procedures were performed as previously described ^36^. Heart tissues were fixed in neutral-buffered 4% paraformaldehyde in PBS, rinsed, dehydrated, paraffin-embedded and tissue sections cut at 8 μm. Sections were stained with Harris Hematoxylin (Sigma #MHS32-1L) and Eosin (Sigma #HT110232-1L) or Trichrome stain kit (Masson) (Merck #HT15-1KT). For immunostaining the samples were blocked with 5% horse serum (Thermo Fisher Scientific #16050122) 0.1% Tween 20 in PBS. Primary antibody used were Isolectine B4488 (1/200; Thermo Fisher Scientific #I21411), Ki67 (1/100; Thermo Fisher Scientific #14-5698-82), ERK1/2 (1/100; Cell signaling #4696), pERK1/2 (1/100; Cell signaling [#4376]), secondary antibody Anti-Rat Alexa 647nm (1/200; Thermo Fisher Scientific # A-21247) and with DAPI (1/1000).

### Quantification of transcript expression

Cells were lysed on Trizol (Thermo Fisher Scientific #15596018) and RNAs were extracted using chloroform, isopropanol and ethanol. Total RNAs were then retrotranscribed to cDNA using AffinityScript Multiple Temperature cDNA Synthesis Kit (Agilent #200436) and qPCR were run on QuantStudio 5 according to PowerUp™ SYBR® Green instructions (Thermo Fisher Scientific #A25742). Samples were normalized to TBP as an endogenous housekeeping gene. List of qPCR primer used: Tbp (mouse) CCCCACAACTCTTCCATTCT; GCAGGAGTGATAGGGGTCAT, Nos3 CCTAGAGCACGAGGCACTG; GTTGTACGGGCCTGACATTT, Col1a1: CTGGATGCCATCAAAGTCT; TCTTGTCCTTGGGGTTCTTG, Col1a2: CTGGTAGTCGTGGTGCAAGT; AATGTTGCCAGGCTCTCCTC, Ki67: TAACCATCATTGACCGCTCC; GGCCCTTGGCATACACAAAA, Col5a1 CTATGACCCGTACTTTGACCC; TCTCCCTTTTGCCCTTTCTC, Col6a2 GTACCCAGGCATCTTCTCCA; ATGGAGGACCCCCAAGAGT, Col14 GAAGCGTGGTTATATCAGAGGG; GGTAGTTTCCTCAGTGATGGTG, Tgf-β TACGCCTGAGTGGCTGTCTTTT; ACAAGAGCAGTGAGCGCTGAAT, Pbp (rat) ACCCCACAACTCTTCCATTC; GGGTCATAGGAGTCATTGGTG, Egr1 GTTGCCTCCCATCACCTATAC; GATGAAGAGGTTGGAGGGTTG, Klf2: CAAGACCTACACC AAGAGTTCG; CCAGTTGCAATGATAAGGCTTC, Biglycan: TTACTGACCGCCTGGCCATCCA; TGCTTAGGA GTCAGGGGG AAGCTGT, Versican: GCTGCCCCGAGCCTTTCTGG; GCGCTTGGCCACAGCACCTC, Decorin: GGTGT CAGCTGGATG CGCTCAC;TGCAGCCCAGGCAAAAGGGTT.

### Cell culture

Cos7 cells were maintained in a DMEM media with 10% of serum (Thermo Fisher Scientific #61965) and 1% of antibiotic antimitotic mix (Gibco #15240062). For the isolation of valvular cells, dissected rat aortic valves were incubated with collagenase 425U/ml (Thermo Fisher Scientific #17101015) during 5 minutes at 37°C, 5% CO2. Valves were scratched to free the cells, centrifuged and plate on a collagen coated plate in the Promocell Endothelial Cell Basal Medium MV completed with the Growth Medium MV SupplementPack (Promocell # C-22120). From that point cells were cultured on collagen functionalized plate for maximum seven passages. For activation of PIEZO on static cell we used Yoda1 (Merck-Sigma SML1558) at 5µM. The calcium imaging was performed using F14201 from Thermo Fisher Scientific.

### Bioreactor

This fluid activation device is adapted to mimic the rapid changes in flow direction. It can master pulsatility, accelerations, decelerations and fast variations of flowrate generating pathophysiological flow waveforms ^29^. 12 hours prior to the WSS stimulation cells were seed (100 000/well) on a 3D collagen matric (rat tail collagen [10 mg/mL concentration, BD Biosciences], 0.1M NaOH and Dulbecco’s Modified Eagle’s Medium 160 [DMEM, Thermo Fisher Scientific #61965 with 10% fetal bovine serum from Thermo Fisher Scientific # A5256701]). The program used mimic the WSS at the tip of the fibrosa side on the aortic valve (max WSS: + 13.69 dyn/cm^2^, min WSS: −11.86 dyn/cm^2^, ΔWSS: 25.56 dyn/cm^2^). The different treatments used were add on the cells 1 hour prior to WSS stimulation: AG1478 at 10 µM (Sigma, #T4182); U0126 at 10 µM (Abcam # 218601-62-8); hEGF 10 mM (Sigma #E9644); GsMTx4 at 10 µM (Sigma # SML3140); PP2 at 200 µM (Sigma #P0042-5MG); SB 202190 at 10µM (Abcam, #ab120638).

### Plasmid reporter

The 265 pb upstream of *Nos3* has been cloned in a plasmid containing the Firefly luciferase gene. A mutated version of the promoter has been generated using the Phusion TM site-directed mutagenesis (Thermo Fisher Scientific) and the following primers: 5’-TCCCATTGTGTGATTTACAGGGGC-3’; 5’-GACAGGAACAAGGCCGGCAGG-3’) previously published ^37^.

### Luciferase assay

Cells were seed at 100 000 cells per well in a 24 wells plate. Transfection in Cos7 cell were done using Lipofectamine (Thermo Fisher Scientific # L3000008). Increase concentration of EGR1 coding plasmid (with a CMV promoter) was transfected with 10 ng of constitutive Renilla luciferase coding plasmid (for normalization), 200 ng of Firefly luciferase reporter and salmon sperm ADN to reach an equivalent quantity of DNA in all conditions. The assay was performed 24 hours after transfection using Dual luciferase assay kit for Promega.

### Adenovirus

Cells were infected with 2X10^6^ plaques forming unit for 100 000 cells. Adenovirus inducing the expression of EGR1 (#ADV-207671) and the GFP (control; #1060) have been order at Vector Biolabs. Cells are harvested 24 hours after infection.

### siRNA

siRNA have been ordered at Ambion for Thermo Fisher Scientific (CCCAACAGUGGCAACACUUtt; AAGUGUUGCCACUGUUGGGtg) and compared again a control siRNA with a Cy3. When the cells reach 80% of confluence 5pmol/µl of siRNA is transfected with 1.5 µl of lipofectamine (Thermo Fisher Scientific). Cells are harvested 24 hours after transfection.

### Western blot

For protein extraction from the collagen gel, pretreatment with type II collagenase (100U/ml; Thermo fisher scientific) at 37°C was performed. Cells were lysed with RIPA buffer (Thermo Fisher Scientific #L3000008) with protease and phosphatase inhibitor (Thermo Fisher Scientific #78440) for 20 minutes on ice. Samples were than hearted at 95°C for five minutes. Sample were migrated on an acrylamide gel (NoverTM WedgeWellTM 4 to 20% (Fisher Scientific #15466814) and transferred on a PVDF membrane. The following antibody were used for protein detection: α-Vinculine (Sigma V9264), α-ERK1/2 (Cell signaling #4696), α-pERK 1/2 (Cell signaling #4376), α-mouse (Thermo Fisher A16011), α-rabbit (Thermo Fisher A16096).

### Zebrafish NO staining

Zebrafish were maintained under standardized conditions ^38^ and in accordance with the European Communities directive 2010/63/EU. Morpholino oligonucleotides (MO) *tnnt2* was obtained from Gene Tools (Philomath, OR, USA) and injected into one-cell stage embryos. The sequence of the injected MO is the following: tnnt2 MO: 5′-CATGTTTGCTCTGATCTGACACGCA3’-3ng. To reveal the presence of NO, larvae were incubated in E3 medium containing 5 mM DAF-FM DA (Life Technologies, D23842), for 1 hour in the dark at 28°C. Fluorescence intensities of the OFT were measured using ImageJ software. Pharmacological treatment: BDM (10 or 15 mM, Sigma-B0733), GsMTx4 (1µM, Smartox Biotechnology), AG1478 (10 µM, Sigma-T4182), SB202190 (10 µM, Abcam ab120638) and PP2 (200 µM, Sigma-P0042) were added to the embryo medium 2 hours (BDM), 4 hours (GsMTx4) and one hour (AG1478, SB202190, PP2) before analysis.

### Cell counts, area measurement and statistical analysis

The nuclei of cells were manually counted in the aortic valve leaflet at E10.5 and E13.5. Immunofluorescent in which DAPI used for total cell counts, or Ki67 for proliferative cell counts, from each of the three discrete primordia. Measurement of the valve aera was quantified using a Zeiss Apotome microscope with Zen software. Image J software was used to measure the surface of the valve and the number of nuclei labeled by DAPI. At least three independent measurements were taken per leaflet, of ten different sections. The values were averaged. Three to six animals were used per genotype for statistical analysis. All sections were 8μm in thickness and covered distal, mid, and proximal regions of aortic valve. The data was presented as total cell counts for each leaflet or for all three leaflets together. Student’s (two-tailed unpaired) t-test, a one-way ANOVA, or two-way ANOVA test were performed using Graphpad Prism 10.

## Results

### Loss of Egfr in valvular cells induces aortic valve hypertrophy

A previous study has shown that, depending on the genetic background, a hypoactive allele of *Egfr* can induce general cardiac hypertrophy and also affect the aortic valves of mouse embryos ^18^. To further explore the mechanism underlying aortic valve enlargement, we specifically deleted the *Egfr* gene in valvular endothelial or mesenchymal (also named interstitial) cells, using the *Egfr^flox^*allele in combination with *Tie2-Cre* ^34^ or *Sm22α-Cre* ^35^, respectively. At three months, histological analysis showed that aortic valve leaflets were thicker in both conditional mutants compared to control (wild-type) embryos (Figure 1 A-D). Morphometric analysis revealed a significant increase in aortic valve leaflets size in *Sm22α-Cre;Egfr^f/f^* mice compared to controls. In contrast, *Tie2-Cre*;*Egfr^f/f^* mice did not exhibit a statistically significant increase in valve size, although some individuals displayed aortic valve hypertrophy, with leaflet dimension ranging from two to four times the average control size observed in controls (Figure 1E).

**Figure 1:**
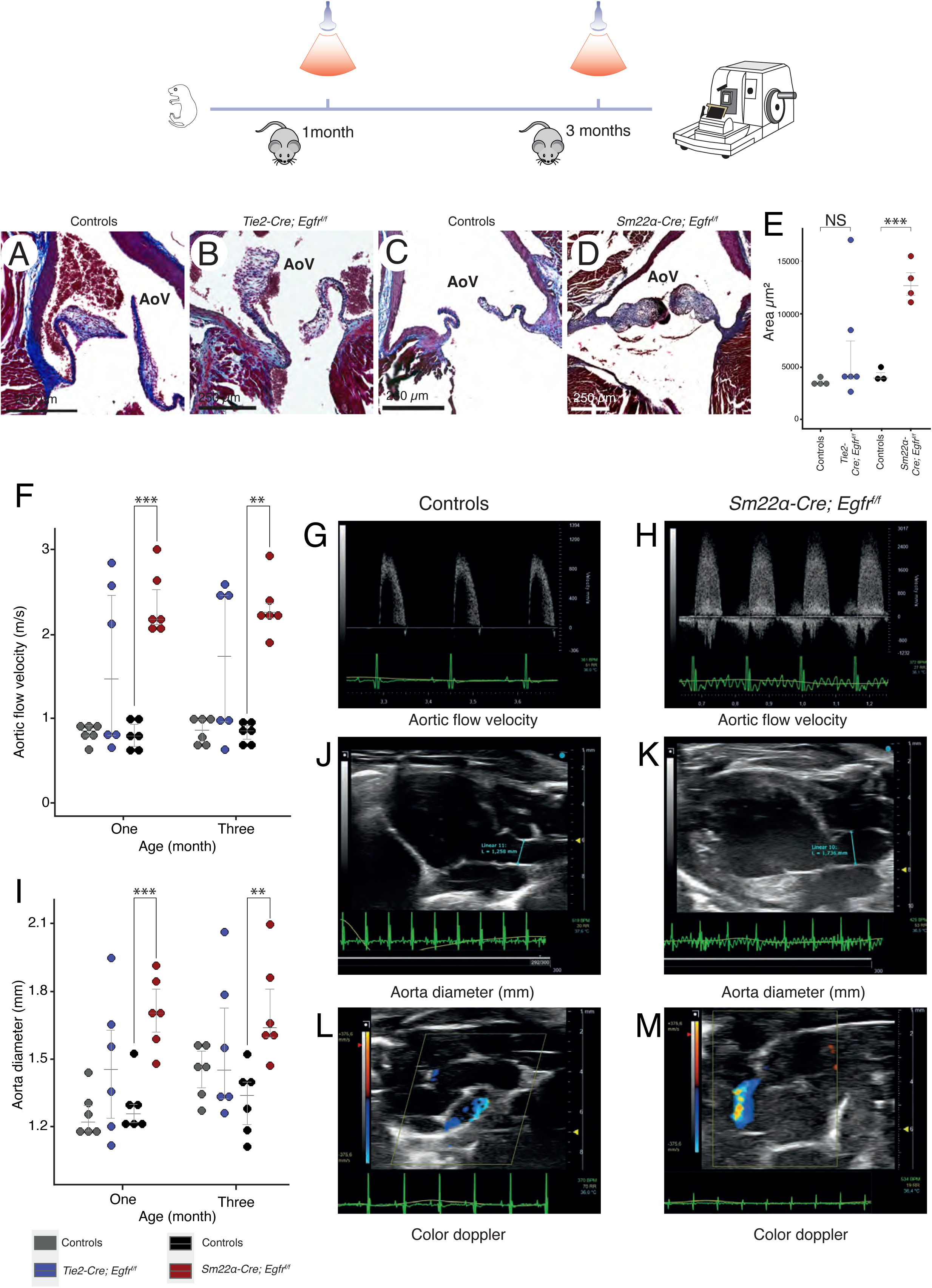
Deletion of the *Egfr* gene in valvular cells induces aortic valve hypertrophy. (**A-D**) Masson’s trichrome staining of three months old mice *Tie2-Cre; Egfr^f/f^*, *Sm22α-Cre; Egfr^f/f^* and controls (Wild-type). (**E**) Quantification of the thickness confirms an enlargement of mutant aortic valves compared to controls (n=3 and 6). The quantification of the valve morphology was performed on the entire valve, not only on one representative slide to avoid any section bias. (**F-H**) Echocardiographic aortic flow velocity (m/s) in controls, *Tie2-Cre;Egfr^f/f^,* and *Sm22α-Cre;Egfr^f/f^* mice (n=6). (**I-K**) Aorta diameter (mm) observed by echocardiography of controls, *Tie2-Cre;Egfr ^f/f^*, and *Sm22α-Cre;Egfr ^f/f^* mice (n=6). (**L,M**) Pulse-wave Doppler recording displays a severe aortic insufficiency (aortic regurgitation) in *Sm22α-Cre;Egfr^f/f^* mice. Controls were obtained from the same litter than the mutants, ensuring that each mutant group had its own corresponding control group. For normally distributed data, the statistical analysis was performed using a pairwise *t*-test, while non-normally distributed data were analyzed using a Wilcox test with Holm correction for multiple testing (NS = p value > 0.05; * = 0.05 < p value > 0.01; ** = 0.01 < p value > 0.001; *** = 0.001 < p value).

To evaluate valve function, we performed echocardiography on conditional *Egfr-cKO* mutant mice at 1 and 3 months of age. Pulse-wave Doppler analysis showed that half of the *Tie2-Cre;Egfr^f/f^*mice had an acceleration of aortic velocity compared to control mice (Figure 1F). Interestingly, animals with accelerated aortic velocity also exhibited aortic valve hypertrophy. Consistent with their morphometric valve defects, *Sm22α-Cre;Egfr^f/f^*mice exhibited significant increased acceleration of aortic velocity (Figure 1F-H). Indeed, hemodynamic evaluation demonstrated a significant increase in aortic flow velocity (805±169 mm/s vs. 2353±381 mm/s; p<3.8*10^-6^) in *Sm22α-Cre;Egfr^f/f^*mice (n=6). Interestingly, mice with accelerated aortic velocity also displayed a dilated aorta (1.296±0.120 mm vs. 1.707±0.158 mm; p<0.00048) (Figure 1I-K). Furthermore, regurgitation was detected in 4 out of 6 *Sm22α-Cre;Egfr^f/f^*mice, compared to 2 out of 6 *Tie2-Cre;Egfr^f/f^* mice (Figure 1L,M). After three months of age, *Sm22α-Cre;Egfr^f/f^* mice also exhibited a significant reduction of fractional shortening between diastole and systole (Supplementary Figure 1). These findings suggest an increase in aortic stroke volume and/or obstruction across the aortic valve due to thickened leaflets. Overall, the data demonstrate that deletion of the *Egfr* gene in valvular mesenchymal cells (*Sm22α-Cre* lineage) or in endothelial cells and their derivatives (*Tie2-Cre* lineage) leads to enlargement of the aortic valve leaflets, resulting in aortic valve dysfunction.

### Aortic valve hypertrophy is caused by excessive proliferation of valvular interstitial cells

To gain deeper insights into the molecular mechanisms underlying aortic valve enlargement in *Egfr-cKO* mutant mice, we performed anatomical and cellular analyses of the aortic valve during development. These analyses focused exclusively on *Sm22α-Cre;Egfr^f/f^* mice due to their fully penetrant phenotype. At E13.5 and E18.5, *Sm22α-Cre;Egfr^f/f^*embryos exhibited enlarged aortic valve leaflets (Figure 2A-E). Valve hypertrophy, evident at E13.5, became more pronounced by E18.5. To examine the transition between the proliferation and maturation phases of valve development, we analyzed the number and proliferative state of valvular mesenchymal cells at E10.5 and E13.5 (Figure 2F,G). Between these stages, the number of mesenchymal cells increased 1.5-fold in controls but doubled in *Sm22α-Cre;Egfr^f/f^* embryos. Correspondingly, the number of proliferative cells (Ki67+) remained constant in controls but doubled in *Egfr* conditional mutant embryos. We further validated these findings using qRT-PCR analysis of micro-dissected OFT regions (Figure 2H), which revealed a 2-fold increase in *Ki67* mRNA expression in *Sm22α-Cre;Egfr^f/f^* embryos compared to controls (Figure 2I). Together, these findings suggest that the enlargement of aortic valve leaflets in *Sm22α-Cre;Egfr^f/f^*mutant embryos is driven by an increased number of valvular mesenchymal cells within the leaflets.

**Figure 2:**
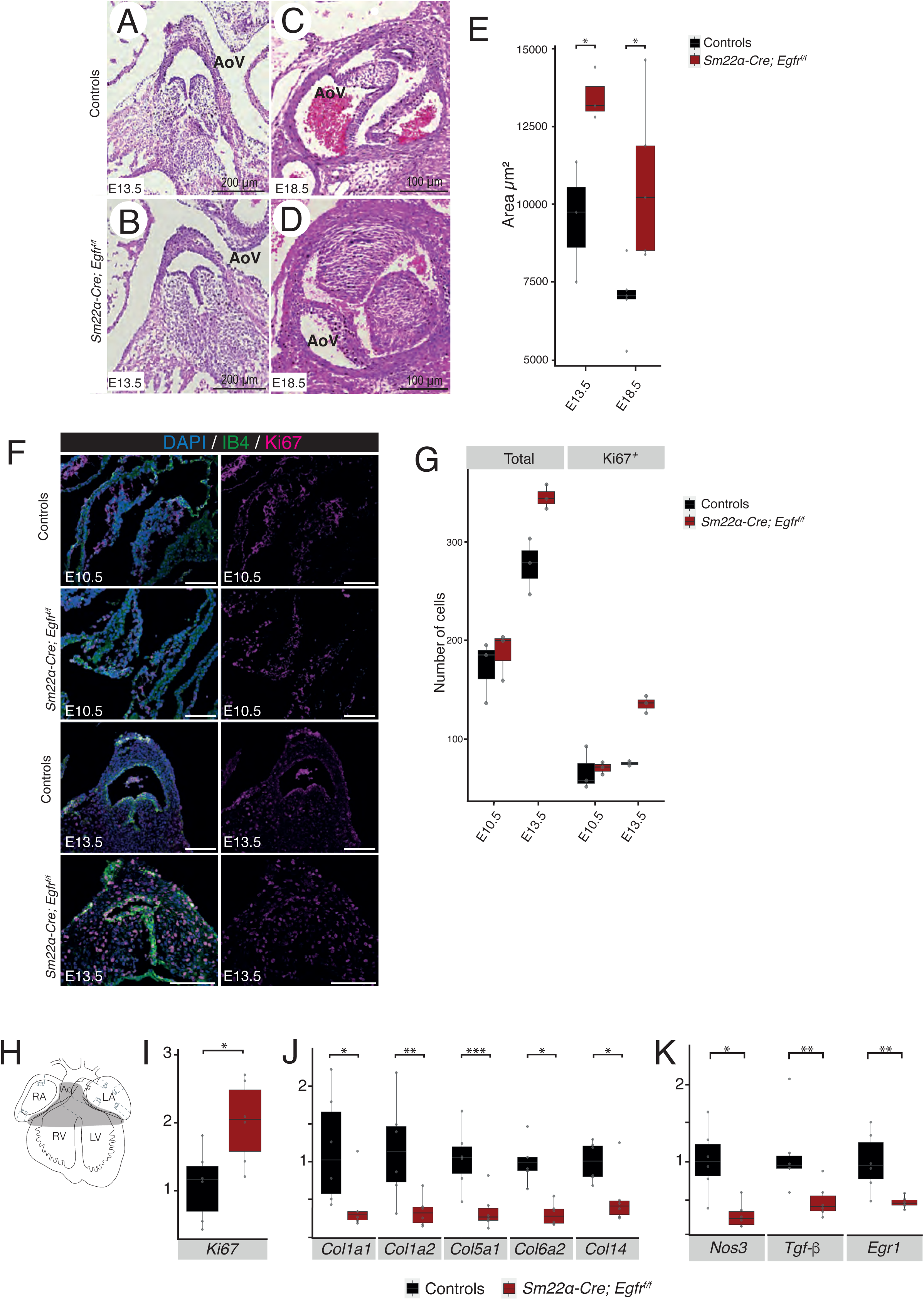
Aortic valve hypertrophy observed in *Egfr*-cKO mutant is caused by excessive proliferation of valvular cells. (**A-D**) Section of E13.5 and E18.5 controls (wild-type) (**A,C**), and *Sm22α-Cre;Egfr^f/f^* (**B**,**D**) hearts (n=3) stained with hematoxylin and eosin. (**E**) Quantification of the thickness confirms an enlargement of mutant aortic valves compared to controls. The quantification of the valve morphology was performed on the entire valve, not only on one representative slide to avoid any section bias. (**F**) Immunolabeling of E10.5 and E13.5 aortic valve against IB4 and Ki67, the nucleus is labeled with DAPI. (**G**) Quantification of the number of cells and the number of proliferative cells (Ki67) at E10.5 and E13.5 of the developing aortic valve (n=3). (**H**) Illustration of the microdissection performed on E13.5 heart for qPCR analysis. (**I-K**) Quantification of mRNA in microdissection E13.5 outflow region (n=6). For normally distributed data, the statistical analysis was performed using a pairwise *t*-test, while non-normally distributed data were analyzed using a Wilcox test with Holm correction for multiple testing (NS = p value > 0.05; * = 0.05 < p value > 0.01; ** = 0.01 < p value > 0.001; *** = 0.001 < p value).

During valve development, differentiated valvular mesenchymal cells express various ECM components, including collagen, elastin, and proteoglycans. qRT-PCR analysis of OFT tissues at E13.5 revealed that several collagen genes were downregulated in *Sm22α-Cre;Egfr^f/f^*embryos compared to controls (Figure 2J). Although, *Biglycan*, *Decorin*, and *Versican* were also examined, their expression levels showed no significant differences (Figure S2). Interestingly, the downregulation of other valve differentiation markers, such as *Nos3, Tgf-β,* and *Egr1*, highlighted an imbalance between the proliferation and differentiation of valvular mesenchymal cells in *Sm22α-Cre;Egfr^f/f^*mutant embryos (Figure 2K). The downregulation of *Egr1* is particularly noteworthy, as it encodes a transcription factor activated by mechanical stress ^39^. In epidermal cells, EGFR has been link to stretch response ^40^. Therefore, we focused our work on the relationship between *Egr1* and *Egfr* to better understand the response of valvular cells to the WSS.

### *Egr1* is able to regulate the expression of *Nos3, Tgf-β*, and *Col1a1*

*Egr1* has previously been linked to cardiovascular pathology including cardiac hypertrophy ^41^, atherosclerosis ^42^ and angiogenesis ^43^. We have recently shown that *Egr1* expression is regulated by WSS in valvular endothelial cells ^29^. To analyze these functional relationships, primary cultures of valvular endothelial cells were isolated from rat aortic valve leaflets. We evaluated whether Egr1 can induce *Tgf-β* and *Nos3* expression using adenoviruses to overexpress human *EGR1* mRNA in valvular endothelial cells. Transduction with Ad-h*EGR1* led to a 4-fold increase in human *EGR1* mRNA levels (Figure 3A). This resulted in a significant increase in *Tgf-β* and *Nos3* expression compared with controls (Ad-eGFP; Figure 3A), indicating that Egr1 regulates the expression of *Tgf-β* and *Nos3*. Next, we used a small interfering RNA (siRNA) against *Egr1* mRNA to knock-down its expression. Valvular endothelial cells transfected with *Egr1* siRNA showed a 75% reduction in *Egr1* expression compared to cells transfected with a commercially available control siRNA (Figure 3B). Furthermore, transfected cells also showed a significant decrease in *Tgf-β* expression, confirming that Egr1 regulates *Tgfβ* expression in valvular endothelial cells (Figure 3C). No changes were detected in *Nos3* expression, likely due to its low basal mRNA levels under control conditions (data not shown). Recent studies identified conserved and functional Gata and Krox20/Egr2 binding sites in the murine -1.5-kb *Nos3* promoter ^44,45^. We examined the promoter region of *Nos3* and identified one evolutionary conserved Egr1-binding site in the proximal region. Luciferase assays demonstrated that Egr1 overexpression in Cos7 cells increased the transcriptional activity of the -265bp region up to 3.5-fold in a dose-response manner (Figure 3D). Transcriptional activity was abolished when the Egr1-binding site was mutated, indicating the importance of this site (Figure 3D). In other physiological contexts, Egr1 is involved in type I collagen production by directly regulating *Col1a1* and *Col1a2* gene transcription ^46^. Similar to *Nos3,* luciferase assays confirmed that Egr1 trans-activates the mouse *Col1a1* promoter (Figure 3E). Together, these results indicate that Egr1 act as an activator of *Tgf-β*, *Nos3*, and *Col1a1*, and suggest that the reduced expression of these genes in *Sm22α-Cre;Egfr^f/f^*mice may result from the transcriptional downregulation of Egr1.

**Figure 3:**
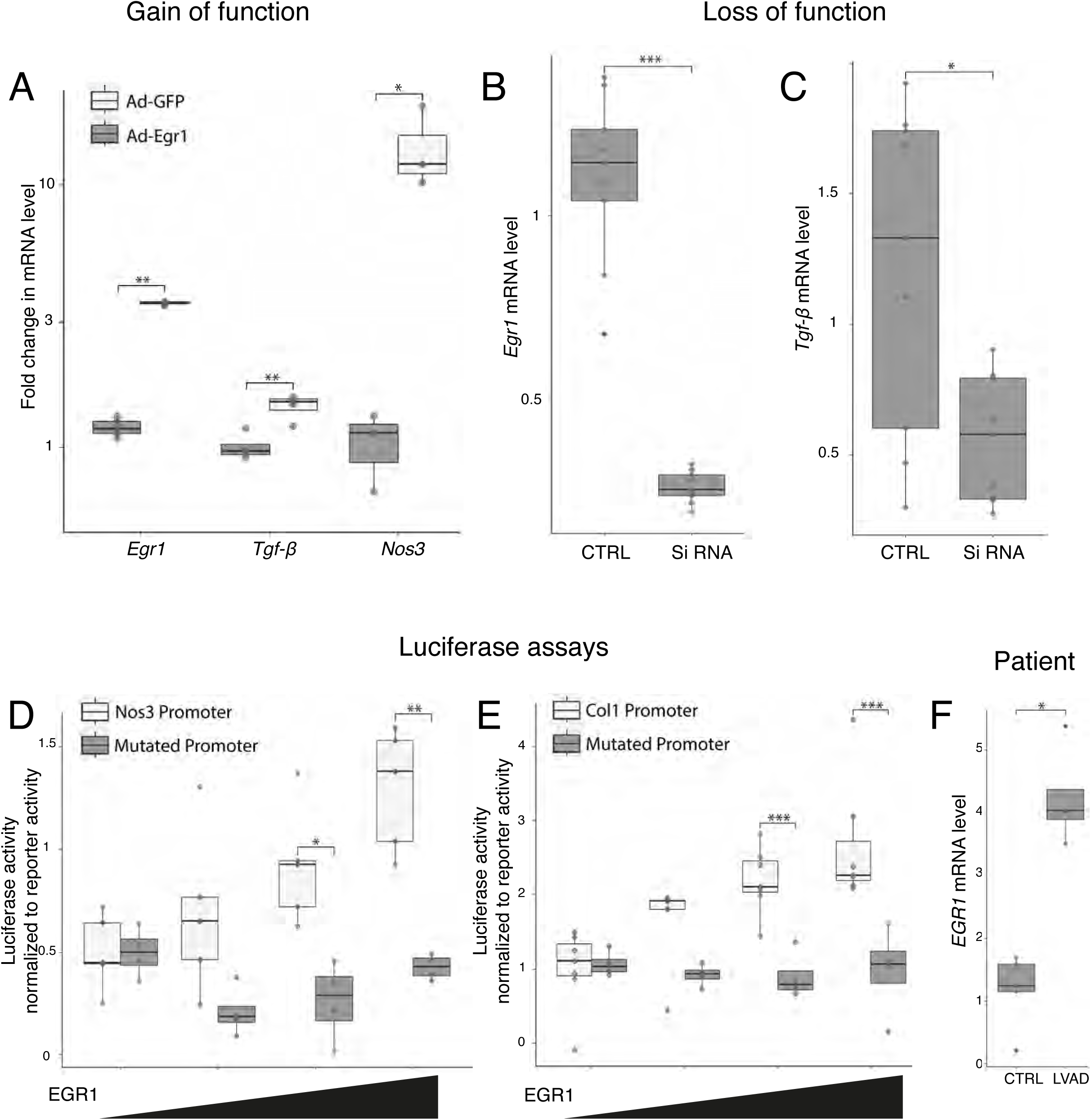
Egr1 regulates the expression of *Nos3*, *Tgf-β*, and *Col1a1*. (**A**) qPCR on valvular endothelial cells infected with adenovirus that induce the expression of Egr1 or the GFP (control) (n=3). (**B**,**C**) qPCR on primary cultured of aortic valve cell treated with Si RNA against *Egr1* or control Si RNA (n=8). (**D**) Luciferase assay on Cos7 transfected cell, the reporter contained a sequence upstream of *Nos3* (this sequence contains one EGR1 binding site which was mutated to confirm its importance; n=4). (**E**) Luciferase assay on Cos7 transfected cell, the reporter contained the sequence upstream of *Col1a1* (this sequence contains one EGR1 binding site which was mutated to confirm its importance) (n=5). (**F**) qPCR on human valves from controls and left ventricular assist device (LVAD) patients to quantify *Egr1* mRNA level (n=4). For normally distributed data, the statistical analysis was performed using a pairwise *t*-test, while non-normally distributed data were analyzed using a Wilcox test with Holm correction for multiple testing (NS = p value > 0.05; * = 0.05 < p value > 0.01; ** = 0.01 < p value > 0.001; *** = 0.001 < p value).

These data highlight Egr1 as a key transcription factor essential for maintaining valve homeostasis. To explore whether *EGR1* expression is altered during human aortic valve remodeling, we analyzed valve samples from patients with left ventricular assist device (LVAD). LVAD increases cardiac output by bypassing the left ventricle and directing blood flow straight into the aorta, leading to biomechanical overload in the aorta root and valves ^47^. This overload triggers aortic valve remodeling ^48^. We obtained aortic valves samples from control donors and LVAD patients after transplantation and performed qRT-PCR analysis. This revealed a four-fold increase in *EGR1* expression in LVAD patient valves compared to controls (Figure 3F). These findings suggest that Egr1 acts as a mechanosensitive transcription factor during cardiac valve formation and remodeling.

### Wall shear stress activates the transcription of Egr1 in an Egfr-dependent manner

The role of Egr1 as a potential activator of *Tgf-β*, *Nos3*, and *Col1a1* downstream of Egfr signaling, coupled with its responsiveness to biomechanical stress, makes it a valuable marker for studying the effects of Egfr signaling during biomechanical activation. We recently developed a novel fluid activation device capable of applying physiologically relevant WSS to the surface of the valvular cells ^29^. Using this device, we confirmed that biomechanical stress induces *Egr1* expression, as demonstrated by its upregulation in valvular endothelial cells exposed to WSS for 60 min compared to static conditions (Figure 4A). To further examine the role of Egfr signaling in biomechanical activation, we pretreated valvular endothelial cells with AG1478, a tyrosine kinase inhibitor of the Egfr. Cells exposed to WSS after 30 min of this treatment showed a significant reduction in *Egr1* mRNA levels (Figure 4A), confirming that Egfr signaling contributes to the response to biomechanical stress. A similar observation was made for the *Klf2* gene, which encodes another transcription factor responsive to biomechanical stress (Supplementary Figure 3A).

**Figure 4:**
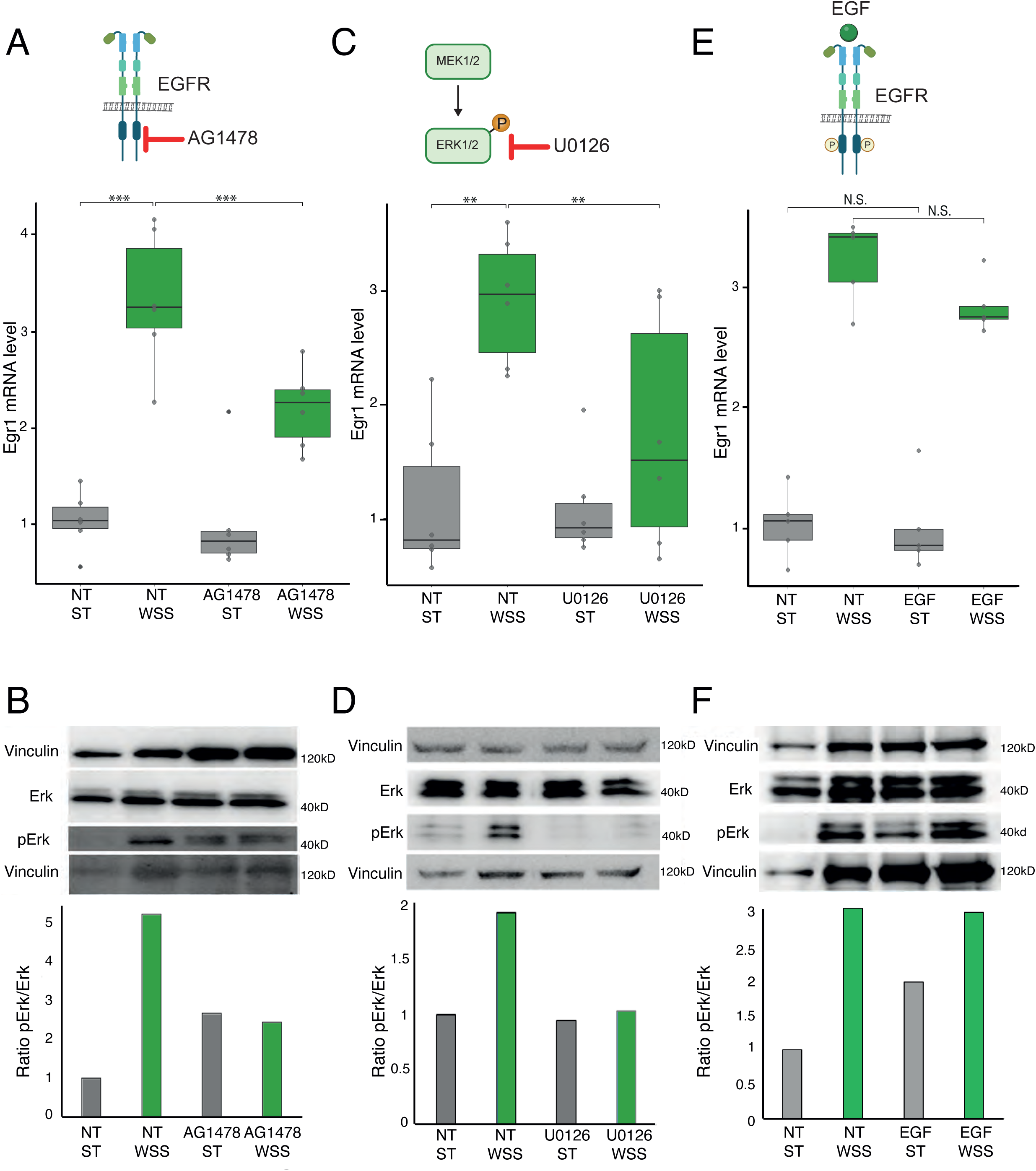
Wall shear stress activates the transcription of *Egr1* in an Egfr-dependent manner. *Egr1* mRNA level in primary cultured of aortic valve cells submitted to WSS and/or AG1478 (**A**, EGFR inhibitor), U0126 (**C**, MEK 1/2 inhibitor), EGF (**E**, EGFR ligand) (n=6). Western-blot analysis of valve cells exposed to WSS and/or AG1478 (**B**, EGFR inhibitor), U0126 (**D**, MEK 1/2 inhibitor), EGF (**F**) with in the lower panel a quantification of the ratio between pERK1/2 and total ERK1/2 expression (n=3). Vinculin was used as internal control. For normal data the statistical analysis has been performed by a pairwise t test and non-normal data were analyzed by a Wilcox test with a holm correction for multiple testing (NS = p value > 0.05; * = 0.05 < p value > 0.01; ** = 0.01 < p value > 0.001; *** = 0.001 < p value). The blots shown in panel D have been spliced.

Both Egfr signaling and Egr1 have been linked to the MAP kinase pathway ^49,50^, which we validated by quantifying the phosphorylation of extracellular signal-regulated kinase (pERK1/2) proteins in valvular endothelial cells exposed to WSS (Figure 4B). Consistent with this, cells pretreated with the AG1478 inhibitor showed reduced pERK1/2 levels after WSS exposure (Figure 4B). To confirm the MAP kinase pathway’s involvement, we used the U0126 inhibitor, which blocks MEK1/2 activation and prevents ERK1/2 phosphorylation (Figure 4C,D). U0126 treatment also consistently inhibited *Egr1* expression in cells exposed to WSS, demonstrating that *Egr1* activation in response to WSS is mediated by the MAP kinase pathway (Figure 4C). We further investigated additional pathways linked to Egfr and *Egr1*. First, the AKT signaling pathway, known to be critical for cardiac development and homeostasis, has been associated with both *Egr1* ^51^ and Egfr ^52^. However, pretreatment with an Akt1/2 kinase inhibitor did not affect *Erg1* expression (Supplementary Figure 3B). Next, we examined the ERK5 pathway, which has been implicated in Egfr-mediated hypertrophy in cardiomyocytes ^53^ and Egr1 activation in intestinal epithelial cells ^54^. Using the BIX02189 inhibitor to block ERK5, we found no effect on *Erg1* upregulation by WSS (Supplementary Figure 3C). Overall, these results emphasize the critical role of the MAP kinase pathway in mediating the response of valvular endothelial cells to WSS.

To further investigate Egfr signaling pathways, valvular cells were treated with recombinant EGF. While EGF enhanced the pERK1/2 to ERK1/2 ratio, it failed to activate *Egr1* expression under static conditions (Figure 4E,D). Similarly, other Egfr ligands were tested but did not induce *Egr1* transcription (Supplementary Figure 3D). These findings suggest that although Egfr signaling is responsive to mechanical shear stress, ligand binding alone is insufficient to fully replicate this activation. These findings suggest that while Egfr signaling is responsive to mechanical shear stress, ligand binding alone cannot fully recapitulate this activation, indicating that the mechanical activation of EGFR-ERK1/2 signaling likely involves distinct cellular mechanosensors that are essential for mediating the effect of WSS.

### PIEZO mechanoreceptor induces a response to WSS through Egfr and the MAP kinase pathway

Our findings revealed that the EGF ligand alone is insufficient to activate *Egr1* transcription in valvular endothelial cells. Previous studies have shown that mechanosensitive Piezo ion channels can respond to shear stress ^55^, and dominant-negative *PIEZO1* mutations have been linked to bicuspid aortic valve defects, implicating a role in valvular cells during cardiac development ^25^. To test whether WSS–dependent Egfr signaling involves Piezo1, valvular endothelial cells were pretreated with GsMTx4, an inhibitor of cationic mechanosensitive channels ^56^. GsMTx4 significantly impaired WSS-induced *Egr1* mRNA activation (Figure 5A). We further explored the role of Piezo1 in biomechanical activation of Egfr signaling by treated cells with the Piezo1 agonist Yoda1, alone or in combination with EGF. Activation of Piezo1 in endothelial valvular cells induces Ca^2+^ influx, as confirmed by Ca^2+^ imaging in Yoda1-treated cells (Supplementary Figure 4A,B). Unlike EGF, Yoda1 alone was sufficient to activate *Egr1* expression in valvular endothelial cells (Figure 5B). Combining EGF with Yoda1 did not enhance *Egr1* transcription compared to Yoda1 alone, although this combination increased ERK1/2 phosphorylation relative to Yoda1 treatment alone (Figure 5C). Furthermore, the U0126 inhibitor, which blocks Yoda1-induced Egr1 activation, confirmed that Yoda1 activates the MAP kinase pathway (Figure 5D) ^57^. Conversely, the AKT inhibitor had no effect (Supplementary Figure 4C). Together, these findings highlight the critical role of the MAP kinase pathway in Piezo1-mediated mechanotransduction, consistent with observations from WSS-stimulated primary valvular cultures.

**Figure 5:**
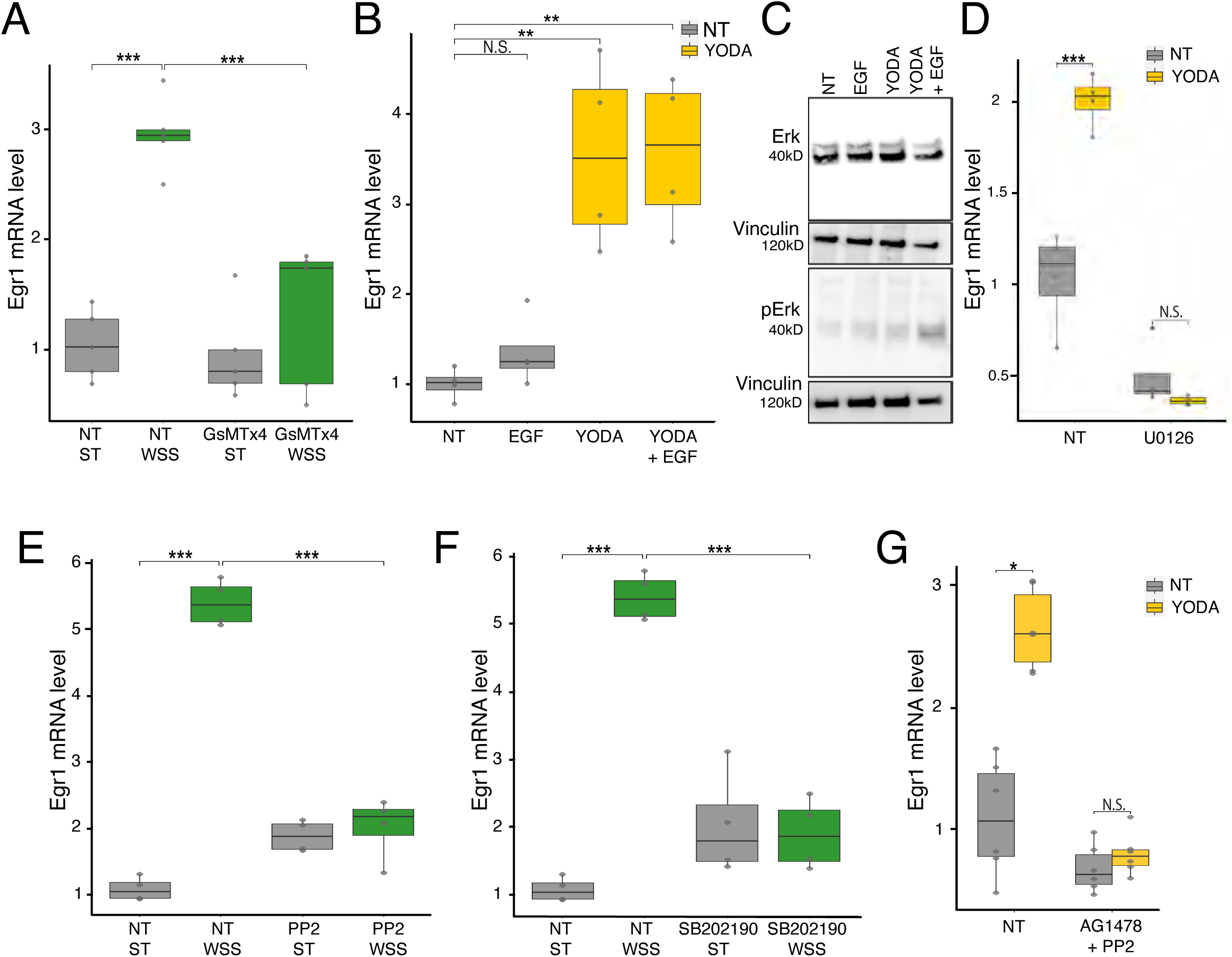
PIEZO mechanoreceptor activates Egfr signaling and the MAP kinase pathway in response to biomechanical stress. (**A,E,F**) *Egr1* mRNA levels in valvular endothelial cells exposed to WSS and/or treated with GsMTx4 (**A**, inhibitor of cationic mechanosensitive channels; n=5), PP2 (**E**, inhibitor for Src-family kinases; n=4), or SB202190 (**F**, p38 MAP kinase inhibitor; n=4). (**B,D,G**) *Egr1* mRNA levels in valvular endothelial cells treated with Yoda1 and/or EGF (**B**), U0126 (**D**, MEK1/2 inhibitor; n=4), or AG1478 + PP2 (**G**, EGFR inhibitor + inhibitor for Src-family kinases; n=6). (**C**) Western-blot analysis of valve cells treated with Yoda1 and/or EGF, showing total ERK1/2, and pERK1/2 expression (n=3). Vinculin was used as internal control. For normally distributed data, the statistical analysis was performed using a pairwise *t*-test, while non-normally distributed data were analyzed using a Wilcox test with Holm correction for multiple testing (NS = p value > 0.05; * = 0.05 < p value > 0.01; ** = 0.01 < p value > 0.001; *** = 0.001 < p value).

To explore the link between Piezo and Egfr signaling, valvular cells were treated with the AG1478 inhibitor prior to Yoda1 stimulation. AG1478 had no effect on Piezo1 activation, suggesting that Egfr-Piezo1 interaction does not involve the tyrosine kinase activity of the Egfr (Supplementary Figure 4D). This aligns with the low minimal increase in *Egf* mRNA observed in valvular endothelial cells exposed to WSS (Supplementary Figure 4E). Piezo1 has been previously linked to Egfr signaling in cancer cells ^58^, where it activates Egfr in a non-canonical manner *via* Src family kinases or p38 MAP kinase. In HeLa cells, Src kinases enable EGFR transduction independently of traditional kinase activity, responding to changes in cell matrix adhesion and Src phosphorylation ^59,60^. Additionally, p38 MAP kinase can phosphorylate Egfr in response to cellular stress ^61^. In valvular endothelial cells, pretreatment with either the Src inhibitor (PP2) or the p38 MAP kinase inhibitor (SB202190) resulted in reduced *Egr1* expression when exposed to WSS, highlighting the role of non-canonical Egfr activation (Figure 5E,F). However, these inhibitors had no effect on Egr1 induction following Piezo1 activation with Yoda1 (Supplementary Figure 4F). Conversely, combining AG1478 (canonical pathway inhibitor) and PP2 (non-canonical inhibitor) effectively blocked Yoda1-induced *Egr1* transcription, suggesting that Yoda1 triggers both canonical and non-canonical Egfr activation (Figure 5G). SB202190 was excluded from these experiments due to its independent enhancement of *Egr1* mRNA levels. Overall, these findings shed light on the complex interplay between mechanical stress from hemodynamic forces and Egfr activation.

### Egfr is implicated in the response to hemodynamic forces in zebrafish OFT

Our data suggests that mechanical forces and Egfr signaling activate Egr1, leading to *Nos3* expression and nitric oxide (NO) production. To validate this *in vivo*, we utilized a NO assay (DAF-FM DA) to assess whether blocking hemodynamic forces or inhibiting Egfr pathways affects NO release in the zebrafish OFT. During zebrafish embryonic development, NO release in the OFT is sustained and coincides with increasing hemodynamic load ^62,63^. In control larvae, a robust NO signal was observed in the developing OFT (Figure 6A,B,K). To test whether NO production depends on hemodynamically forces, we reduced the heart rate by treating zebrafish with 10 mM 2,3-butanedione monoxime (BDM) ^27,64^ which led to a significant decrease in NO production. Complete cessation of the heartbeats with 15 mM BDM abolished the NO signal entirely (Figure 6C,D,K). Similar results were obtained using a *troponin T* (*tnnt2*) morpholino, which abolishes cardiac contractions ^65^. In *tnnt2* morphants, NO production in the OFT was significantly reduced, confirming its dependence on hemodynamic forces (Figure 6E,K). We further investigated mechanosensitive channels by treating larvae with GsMTx4, which significantly reduced NO production in the OFT (Figure 6F,K), implicating a potential role for Piezo1 in detecting these forces. To explore the role of Egfr, we tested inhibitors of canonical (AG1478) and non-canonical (PP2, SB202190) Egfr pathways. AG1478 alone or in combination with PP2 significantly reduced NO production (Figure 6G-J,K). Higher doses or prolonged exposure resulted in lethality, suggesting that Egfr pathways are crucial for overall zebrafish development. The differential response to PP2 and SB202190 may reflect evolutionary divergences in signaling pathways. However, the reduction of NO release by AG1478 supports the role of Egfr in regulating *nos3* expression and NO production. These finding reinforce the importance of Egfr signaling in responding to hemodynamic forces during valve development.

**Figure 6:**
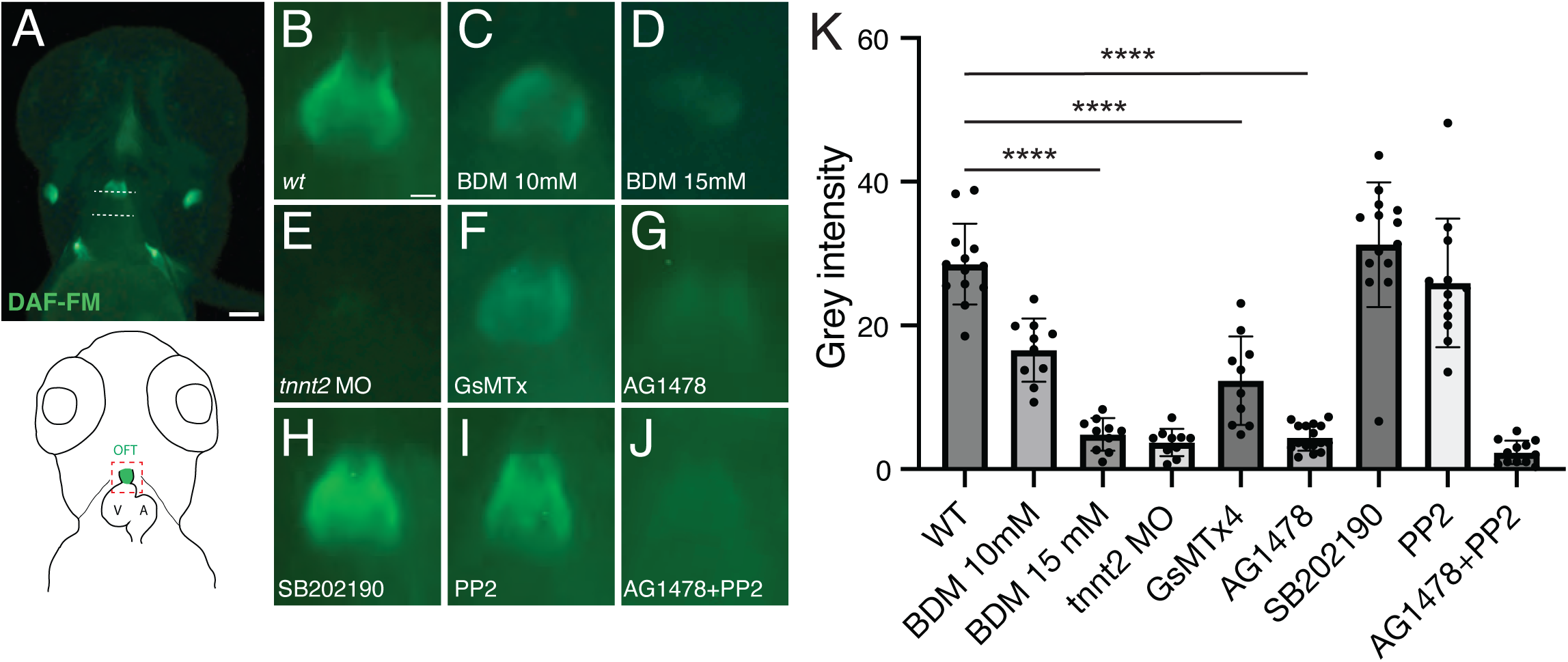
Hemodynamic-dependent nitric oxide production in the zebrafish outflow tract. (**A**) WT zebrafish larvae treated with DAF-FM DA to reveal NO production in the OFT (bisected by upper dashed white line). Dashed white lines indicate the regions where grey scale intensity was measured, the upper line bisects the OFT and the lower line is used to measure the background intensity. (**B-J**) Higher magnification images of the OFT assayed with DAF-FM DA of a DMSO (**B**, n=13), 10mM BDM (**C**, n=10), 15mM BDM (**D**, n=10) treated, *tnnt2* morphant (**E**, n=10), GsMTx4 (**F**, n=10), AG1478 (**G**, n=16), SB202190 (**H**, n=14), PP2 (**I**, n=12I), AG1478+PP2 (**J**, n=12) treated larva. (**K**) Graph showing the mean values of grey scale intensity. Error bars indicate the SEM. One-way ANOVA (**** P<0.0001) followed by student’s unpaired homoscedastic two tailed t-test (****P<0.0001) were used for statistical analysis. Scale bars: 100μm (A); 20μm (B).

### The inability of EGF to induce the transcription of Egr1 appears to be linked to the nuclear translocation of pERK1/2

Our findings emphasize Egfr as a critical mediator of the response to WSS. However, treatment with EGF alone did not increase *Egr1* expression, despite activating the MAP kinase pathway, as evidenced by ERK1/2 phosphorylation (Figure 4E,F). To investigate this unexpected result, we performed immunolabeling on valvular endothelial cells treated with EGF or Yoda1, using antibodies against ERK1/2 and pERK1/2 (Figure 7A). In untreated cells, pERK1/2 levels were very low and distributed between the cytoplasm and nucleus (Figure 7A,B). Yoda1 treatment, however, caused a robust accumulation of pERK1/2 in the nucleus (Figure 7A,B). By contrast, EGF treatment results in a strong pERK1/2 signal surrounding the nucleus but a significantly reduced nuclear presence (Figure 7A,B). Quantification confirmed that EGF induced a much lower proportion of nuclear pERK1/2 compared to Yoda1 treatment (Figure 7C). These findings suggest that, unlike Yoda1, EGF fails to induce the nuclear translocation of pERK1/2 necessary for Egr1 transcription, providing a mechanistic explanation for its inability to activate Egr1 in valvular cells exposed to WSS.

**Figure 7:**
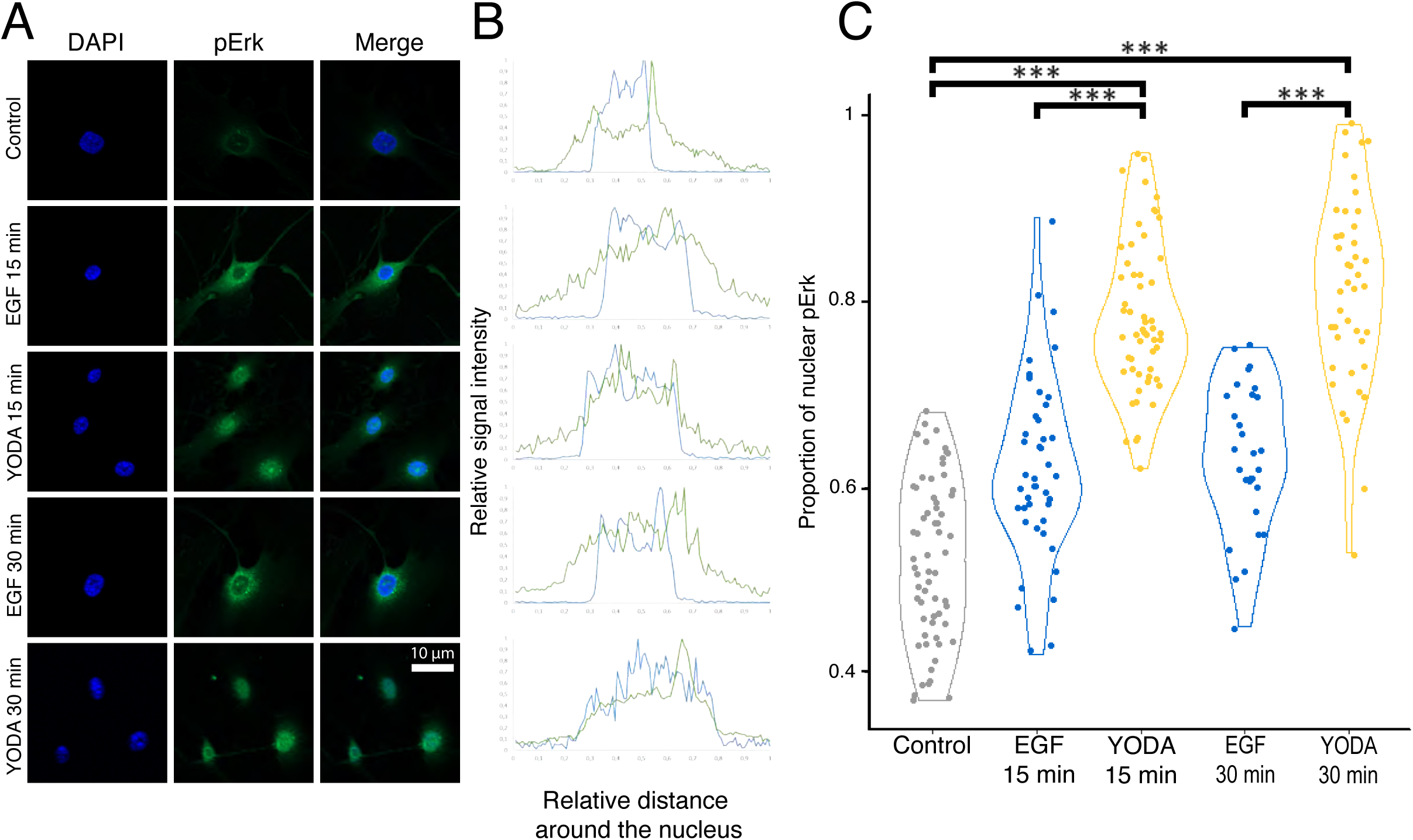
The failure of EGF to stimulate *Egr1* transcription appears to be tied to the nuclear translocation dynamics of pERK1/2. (**A**) Primary cultured of aortic valve cell treated with EGF or Yoda1 immunolabeled of against pERK (nucleus is labeled with DAPI) (n=3). (**B**) Spatial distribution of pERK analyzed with Zeiss Zen software. (**C**) Quantification of pErk cell repartition was performed on ImageJ using the DAPI to define the nuclear region (n=60). For normally distributed data, the statistical analysis was performed using a pairwise *t*-test, while non-normally distributed data were analyzed using a Wilcox test with Holm correction for multiple testing (NS = p value > 0.05; * = 0.05 < p value > 0.01; ** = 0.01 < p value > 0.001; *** = 0.001 < p value).

## Discussion

Aortic valve formation is a complex multistep process that begins by the growth of endocardial cushions in the OFT. This is induced by signals originating from both the myocardium and the endothelium, which control the endothelial-to-mesenchymal transition (EndMT) of endocardial cells in the OFT region. Initiation of the EndMT is followed by the proliferation of the valve primordia and is completed by a phase of refinement and maturation, during which the primitive structures develop into fully functional, anatomically distinct cardiac valves ^3,66^. Proper regulation of these processes is crucial for normal valve development, and disruptions can lead to valve disorders. In this study, we illustrate the key role of Egfr in the transition from proliferation to maturation. The absence of Egfr in both endothelial and mesenchymal cells lead to an extended proliferative stage and delayed maturation, resulting in hypertrophy of the aortic valve leaflets. At postnatal and adult stages, this phenotype results in a stenosis, regurgitation and aortic dilatation. Analysis of the conditional deletion of the *Egfr* gene during valve development revealed that Egfr signaling is crucial for regulating the *Egr1* gene, which encodes a transcription factor that is activated by mechanical stress. This result was confirmed *in vitro* using a unique flow activation device that applies WSS to valvular endothelial cells. Inhibition of Egfr activation with AG1478 reduced the cellular response to WSS, as evidenced by reduced Erk1/2 phosphorylation and decreased *Egr1* mRNA levels. Equivalent inhibition can be achieved by blocking cellular sensitivity to mechanical stretch with GsMTx4, which reduces the transcription of *Egr1*. In Zebrafish, the use of GsMTx4 or AG1478 inhibits the response to hemodynamic forces. Conversely, activating the mechanosensor Piezo1 with Yoda1 induces the transcription of *Erg1* and the phosphorylation of ERK1/2. While treatment with EGF activates the MAP kinase pathway, it does not induce *Erg1* transcription. Other mechanisms of Egfr activation by WSS have also been analyzed. For example, in HeLa cells, the PIEZO can activate the Egfr pathway independently of ligand binding, triggering a cellular response ^58^. This activation is also observed in valvular endothelial cells, where blocking non-canonical pathways decreases the response to WSS. In summary, EGFR is essential for the WSS response in the aortic valves. It is an essential component of a mechanosensitive pathway triggered by PIEZO1, which activates MAP kinase signaling and induces *Egr1* transcription (Figure 8).

**Figure 8:**
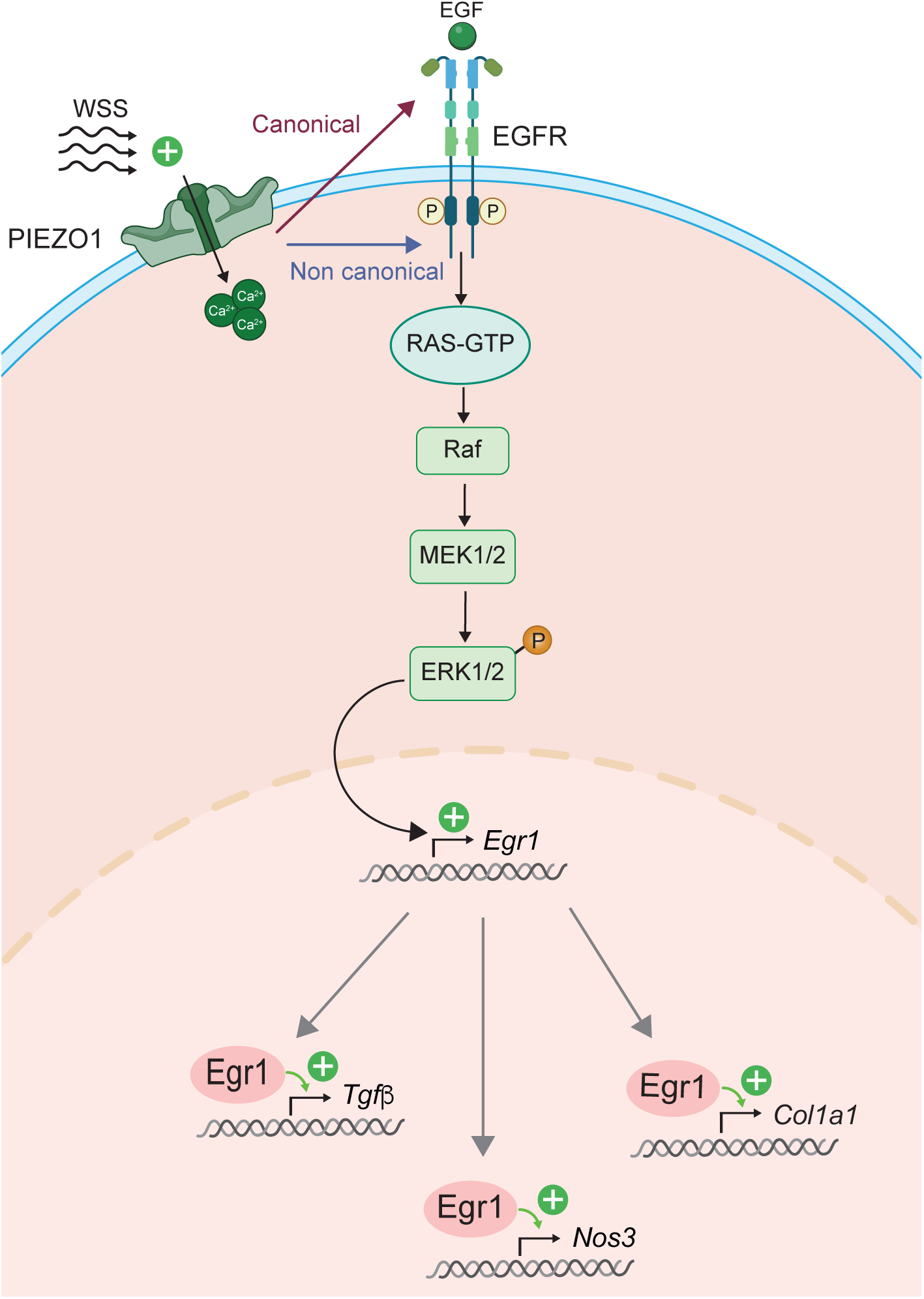
Model of EGFR activation in the WSS Response.

In contrast to WSS stimulation, the canonical inhibitor AG1478 and the non-canonical inhibitors PP2 and SB202168 had no impact on the response to Yoda1, a specific agonist of PIEZO1. To counteract the effects of Yoda1, we used an inhibitor cocktail containing PP2 and AG1478, which blocks both the activation and autophosphorylation of EGFR. Given the distinct responses between WSS and Yoda1, we propose the following hypothesis: the flow generated by our device is designed to mimic the physiological WSS response by activating various pathways in a more controlled and nuanced way. When an element of these pathways is inhibited (as seen with AG1478), the response of valvular cells is diminished. In contrast, a brief treatment with Yoda1 provides a short and intense stimulation of PIEZO1, sufficient to induce a response comparable to one hour of WSS. This brief treatment appears to circumvent the blockade of canonical or non-canonical Egfr activation. However, the combined use of these inhibitors completely abolishes the cellular response. The necessity of employing an inhibitor cocktail alongside Yoda1 treatment to suppress the response underscores the significance of these distinct pathways and their crosstalk. Further investigations are needed to identify the mechanisms involved in the ligand-independent activation of Egfr, which will enhance our understanding of pathway interactions.

In this study, we showed that the MAP kinase pathway is crucial for cellular signaling. While there is crosstalk between canonical and non-canonical activation pathways, blocking ERK1/2 phosphorylation in valvular cells notably reduced their response to both WSS and Yoda1 stimulation. Other inhibitors targeting pathways like AKT and ERK5 were tested but did not significantly impact the response. Our findings suggest that, in valvular endothelial cells, the MAP kinase pathway is the primary signaling mechanism in response to EGF, Yoda1, and WSS stimulation.

EGF, Yoda1, and WSS all induce ERK1/2 phosphorylation, but they differ in their effects on *Egr1* transcription. Unlike EGF, which does not influence Egr1 expression, both Yoda1 treatment and WSS stimulation significantly upregulate Egr1 transcription. Analysis of pERK1/2 localization revealed that Yoda1 treatment promotes its translocation into the nucleus, whereas in EGF-treated cells, pERK1/2 remains perinuclear. This nuclear exclusion of pERK1/2, increasingly recognized in the literature ^67–69^, is being explored as a potential target for cancer therapy ^70^. This mechanism may be critical in environments where mechanical stress maintains homeostasis. Supporting this, a recent study on myocardial cells under stretch conditions identified an alternative ERK1/2 activation pathway via EGFR ^71^. In the context of cardiac valve development, this activation may be essential for transitioning from the proliferation phase to maturation by integrating mechanical stress with EGF signaling. Further studies into the MAP kinase pathway during valve development could provide valuable insights into these processes. Thus, exploring the location of WSS-, EGF-, and Yoda1-induced ERK phosphorylation could shed light on these specific responses.

Egr1 and Tgf-β play critical roles in valve development and are correlated in various contexts, including cardiac hypertrophy ^32^ and pulmonary arterial hypertension ^72^. Notably, *Tgf-β* expression is reported to be downregulated in *Egr1*^-/-^ mice ^73^. In *Sm22α-Cre;Egfr^f/f^* mice, a correlation between *Erg1* and *Tgf-β* expression levels was observed. Similarly, in cultured valvular cells, Egr1 overexpression promoted *Tgf-β* expression, while knockdown via siRNA reduced *Tgf-β* levels. These findings underscore the strong interaction between Egr1 and Tgf-β in valve development and homeostasis. Given the well-established role of Tgf-β in valve pathologies ^74–76^, further understanding of its regulation could pave the way for novel therapeutics approaches.

Here we show that the EGFR-PIEZO1 signaling axis plays a distinct yet interconnected role in valvular cells during valve development and homeostasis. In valvular endothelial cells, EGFR activation is crucial for mediating responses to WSS, primarily through the MAP kinase pathway, leading to the upregulation of *Egr1* and other mechanosensitive genes. PIEZO1, as a mechanoreceptor, facilitates calcium influx in response to mechanical stimuli, augmenting EGFR-mediated signaling. This interaction highlights the importance of mechanical cues in endothelial cell function, integrating physical forces with biochemical signaling to regulate transcriptional responses such as those driven by Egr1. Analysis of conditional knockout models reveal that the absence of Egfr in mesenchymal cells results in prolonged proliferation and valve hypertrophy, underscoring EGFR’s role in transitioning from proliferation to maturation phases. Unlike valvular endothelial cells, where mechanical stress is a primary activator, mesenchymal cell responses to EGFR signaling are more likely influenced by paracrine factors and downstream pathways regulating ECM composition. Therefore, the EGFR-PIEZO1 axis orchestrates a coordinated response across endothelial and mesenchymal cells, ensuring proper valve morphogenesis and functionality by integrating biomechanical and molecular signals. Further investigation into how these pathways interact in a spatial and temporal manner could provide deeper insights into their specific roles in valve biology and pathology.

## Author contribution Statement

Conceptualization: AG, DM, A.F, SZ.; Methodology: AG, DM, AF, CJ, EB, VD, LL, MH, AS.; Investigation: AG, DM, AF, CJ, EB, VD, LL, MH, AS, HAO, AT, JFA, SZ.; Supervision: AF, VD, SZ.; Writing – original draft DM, SZ.

## Confict of interests

The authors have declared that no competing interests exist.

## Acknowledgments

We thank Robert Kelly and Fabienne Lescroart for careful reading of the manuscript. The authors gratefully acknowledge personnel of the animal facility and imaging platforms of Marseille Medical Genetics.

