## Supplementary material for "EGFR-Mediated Mechanotransduction in Aortic Valve Cells: A Key Pathway in Response to Wall Shear Stress"

### Supplementary Materials

**Figure S1: Cardiac function in *Egfr*-cKO mice.** (**A**) The one- and three-month echocardiographic data for fractional shortening during the systole compared to the diastole observed in controls (wild-type), *Tie2-Cre;Egfr*$^{f/f}$, *Sm22α-Cre;Egfr*$^{f/f}$ mice (n=6). (**B,C**) Left ventricular M-mode imaging in controls and *Sm22α-Cre;Egfr*$^{f/f}$ mice for fractional shortening analysis. For normally distributed data, the statistical analysis was performed using a pairwise *t*-test, while non-normally distributed data were analyzed using a Wilcox test with Holm correction for multiple testing (NS = p value > 0.05; * = 0.05 < p value > 0.01; ** = 0.01 < p value > 0.001; *** = 0.001 < p value).

**Figure 2: Quantification of mRNA in microdissection outflow region of E13.5 controls (wild-type) and *Sm22α-Cre;Egfr*$^{f/f}$ hearts.** For normally distributed data, the statistical analysis was performed using a pairwise *t*-test, while non-normally distributed data were analyzed using a Wilcox test with Holm correction for multiple testing (NS = p value > 0.05; * = 0.05 < p value > 0.01; ** = 0.01 < p value > 0.001; *** = 0.001 < p value).

**Figure S3: Wall shear stress activation.** (**A**) *Klf2* mRNA levels in valvular endothelial cells exposed to WSS and/or EGFR inhibitor AG1478 (n=4). (**B,C**) *Egr1* mRNA levels in valvular endothelial cells exposed to WSS and/or AKT kinase inhibitor (**B**) or ERK5 inhibitor (**C**; n=4). (**D**) *Egr1* mRNA levels in cells treated by HB EGF (n=4). For normally distributed data, the statistical analysis was performed using a pairwise *t*-test, while non-normally distributed data were analyzed using a Wilcox test with Holm correction for multiple testing (NS = p value > 0.05; * = 0.05 < p value > 0.01; ** = 0.01 < p value > 0.001; *** = 0.001 < p value).

**Figure S4: Calcium Imaging in valvular endothelial cells.** Egr1 and Egf expression, and signaling pathway inhibition in valvular endothelial cells treated with Yoda1 and WSS. (**A**) Calcium imaging on valvular endothelial cells treated with Yoda1, (**B**) signal quantification with ImageJ. *Egr1* mRNA level in primary cultured of aortic valve cell submitted to Yoda1 and/or AKT inhibitor (**c**; n=4), AG1478 (**D**, EGFR inhibitor; n=4), or PP2 or SB202190 (**F**, inhibitor for Src-family kinases or p38 MAP kinase inhibitor; n=4). (**E**) *Egf* mRNA level in primary cultured of aortic valve cell submitted to WSS (n=6). For normally distributed data, the statistical analysis was performed using a pairwise *t*-test, while non-normally distributed data were analyzed using a Wilcox test with Holm correction for multiple testing (NS = p value > 0.05; * = 0.05 < p value > 0.01; ** = 0.01 < p value > 0.001; *** = 0.001 < p value).

Supplementary Figure 1

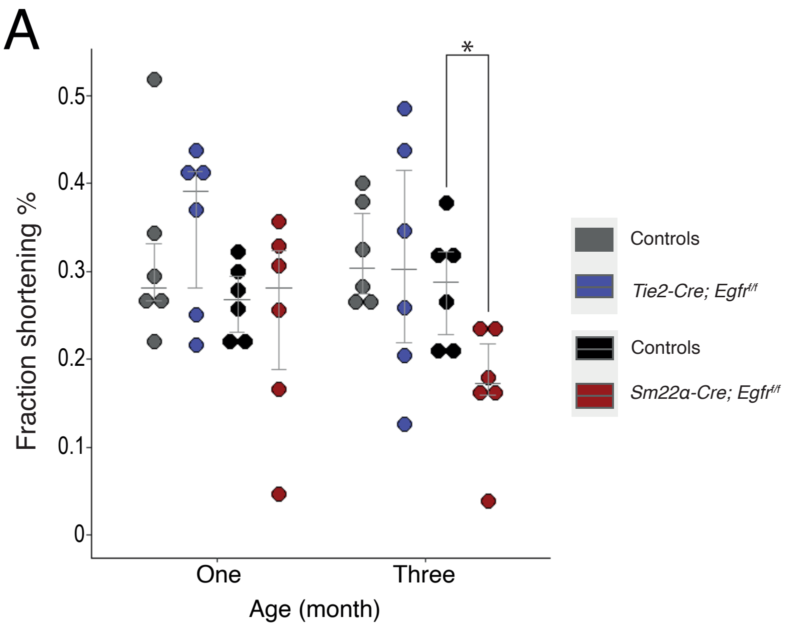


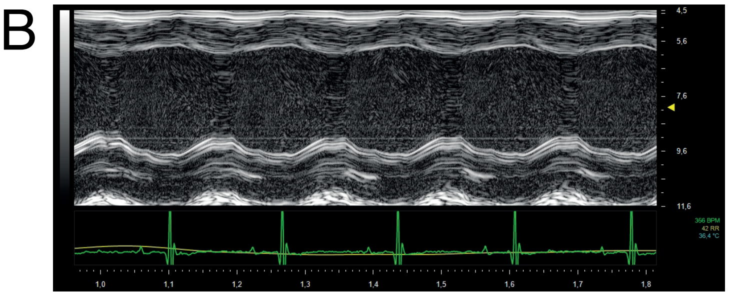


Left ventricular M-mode of Controls

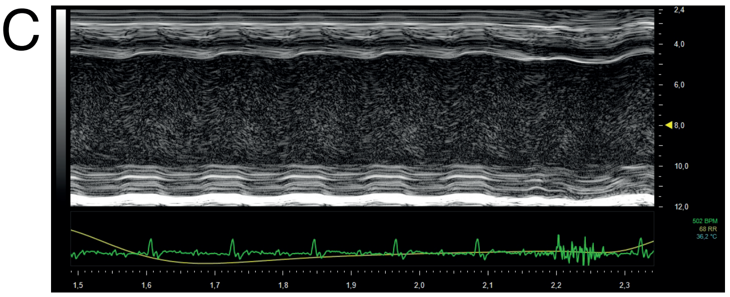


Left ventricular M-mode of *Sm22α-Cre; Egfr*$^{f/f}$

Supplementary Figure 2

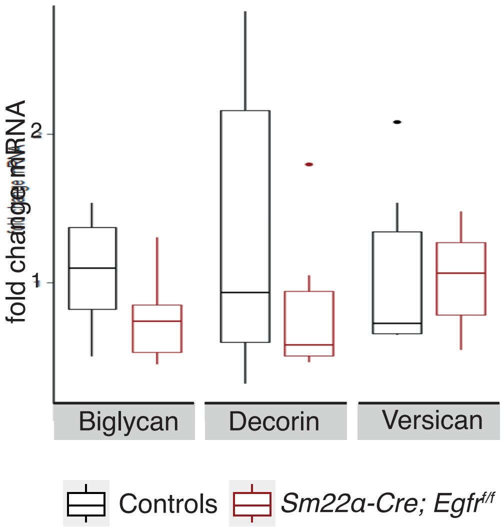

Supplementary Figure 3

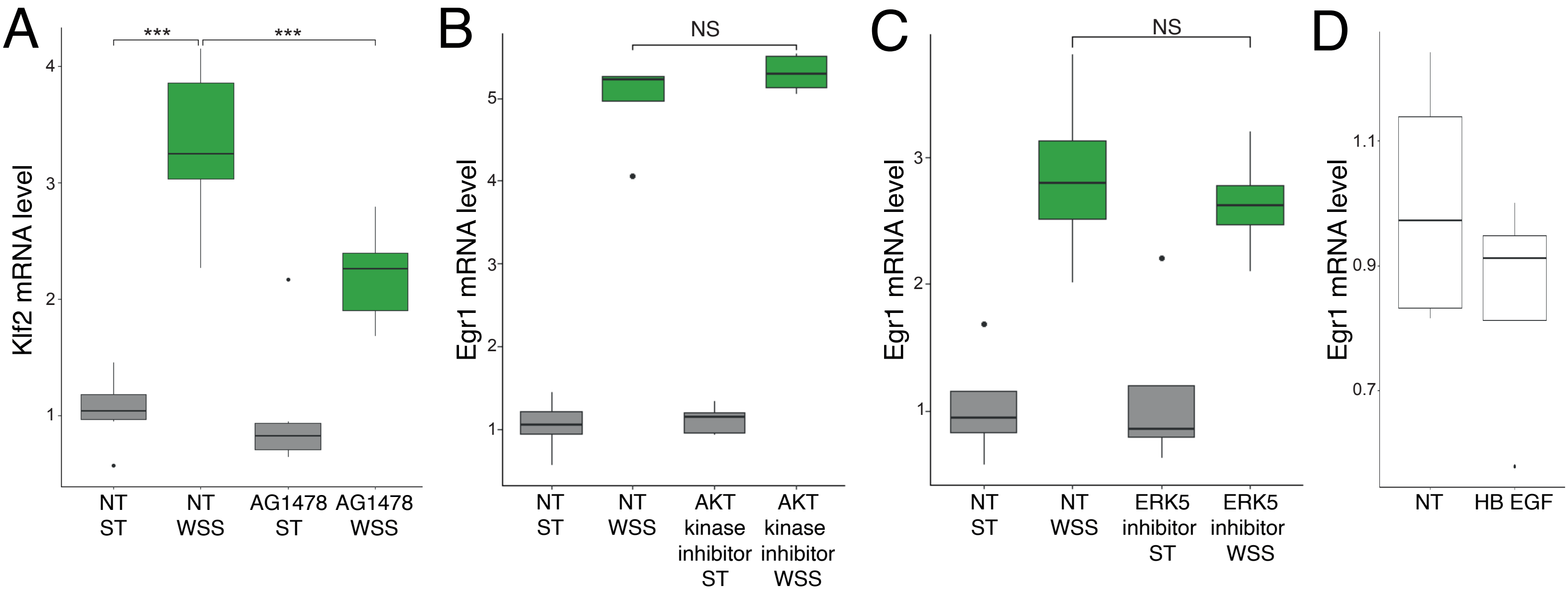

Supplementary Figure 4

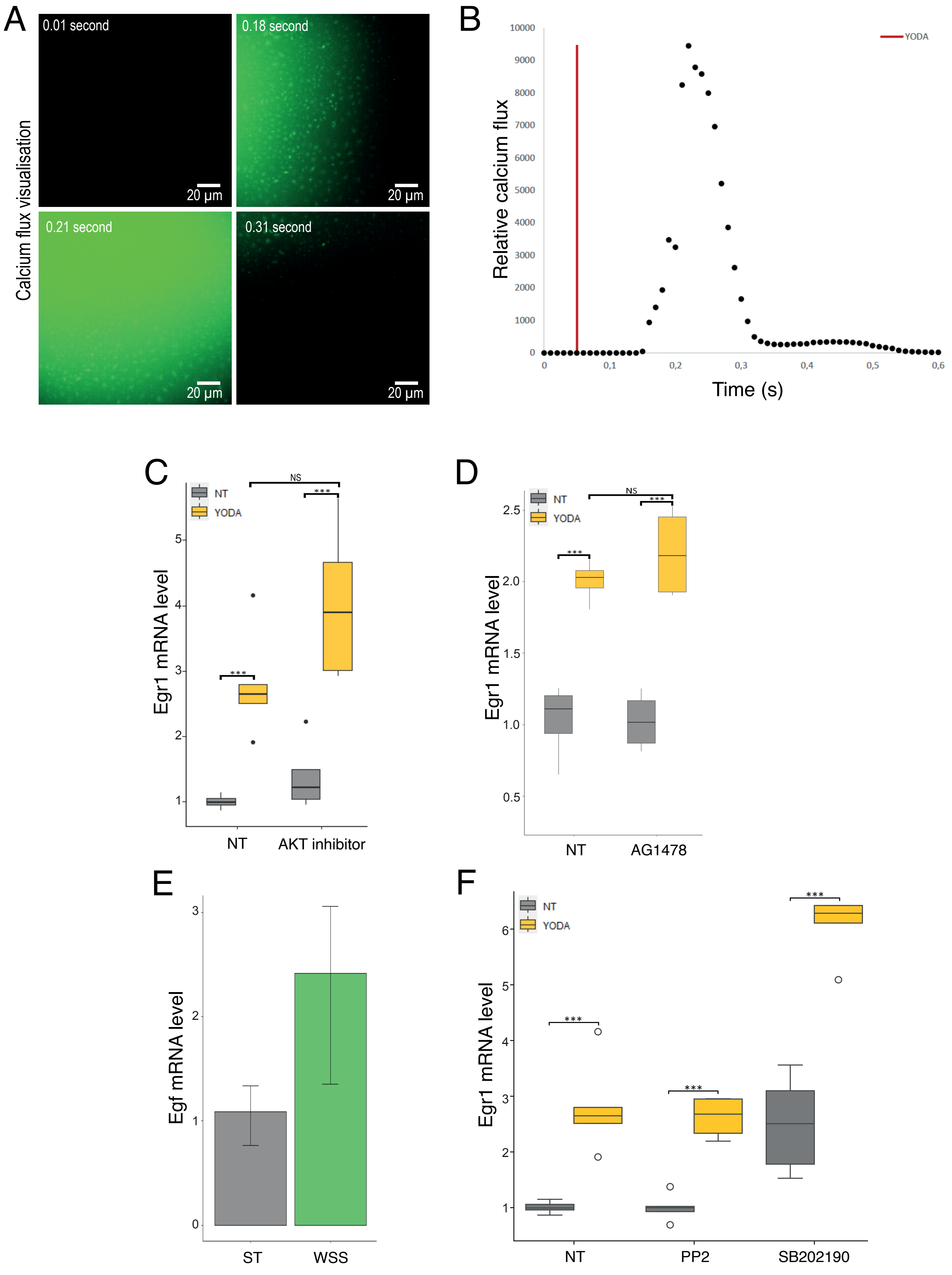
